# Plants export 2-monopalmitin and supply both fatty acyl and glyceryl moieties to arbuscular mycorrhizal fungi

**DOI:** 10.1101/2021.01.19.427311

**Authors:** Leonie H Luginbuehl, Harrie van Erp, Henry Cheeld, Kirankumar S Mysore, Jiangqi Wen, Giles ED Oldroyd, Peter J Eastmond

## Abstract

Arbuscular mycorrhizal fungi (AMF) rely on their host plants to provide them with fatty acids (FA), but the precise form(s) in which they are supplied is still unclear. Here we show that ectopic expression of the transcription factor REQUIRED FOR ARBUSCULAR MYCORRHIZATION 1 (RAM1) can drive secretion of 2-monoacylglycerols (2MGs) from *Medicago truncatula* roots and that their main FA moiety is palmitic acid, although myristic acid and stearic acid were also detected. RAM1-dependent 2MG secretion requires the acyl-acyl carrier protein thioesterase FATM, the glycerol-3-phosphate (G3P) acyltransferase RAM2 and the ATP binding cassette transporter STR. Furthermore, ^14^C glycerol labelling experiments using mycorrhizal *M. truncatula* roots that are deficient in glycerol kinase, FAD-dependent G3P dehydrogenase and the G3P acyltransferase RAM2 suggest that most of the glyceryl moieties in *Rhizophagus irregularis* storage lipids are provided by their host plant through the 2MG pathway. Taken together, our data support the hypothesis that the plant exports 2MGs across the peri-arbuscular membrane in mycorrhizal roots and that the AMF receive and utilise both the FA and glyceryl moieties to make their storage lipids.

## INTRODUCTION

Most plants can form a symbiotic association with arbuscular mycorrhizal fungi (AMF) from the phylum Glomeromycota (Smith and Read, 2008). In this association the plants receive mineral nutrients (mainly phosphate) absorbed by the fungal mycelium that permeates the soil and in return the plants provide organic carbon (Smith and Read, 2008). The transfer of carbon was first demonstrated ∼50 years ago, and until recently it was thought that the bulk of this carbon was supplied as sugars (Smith and Read, 2008). Metabolic tracer studies have established that carbon is transferred to AMF in the form of hexoses (Shachar-Hill et al., 1995; Solaiman and Saito, 1997). However, within the AMF mycelium most of the carbon is stored as lipids (triacylglycerols; TAGs), which are transported throughout the mycelium and accumulate in vesicles and spores (Bago et al., 2002). Initially it was proposed that AMF synthesise fatty acids (FA) in the intra-radicular mycelium, but evidence for this was indirect (Pfeffer et al., 1999; Trépanier et al., 2005). With the completion of the first AMF genome, it became evident that AMF may in fact be FA auxotrophs, because *Rhizophagus irregularis* appears to lack a type-1 FA synthase (Wewer et al., 2014). Subsequently, we (and others) have shown that these obligate biotrophs obtain FA directly from their host plants (Bravo et al., 2017; Jiang et al., 2017; Keymer et al., 2017; Luginbuehl et al., 2017), using a metabolic pathway (Fig. 1) that appears similar to the one that supplies oxygenated FA monomers for the synthesis of plant extracellular lipid polyesters, such as cutin and suberin (Li et al., 2007; Yang et al., 2010).

**Fig. 1.**
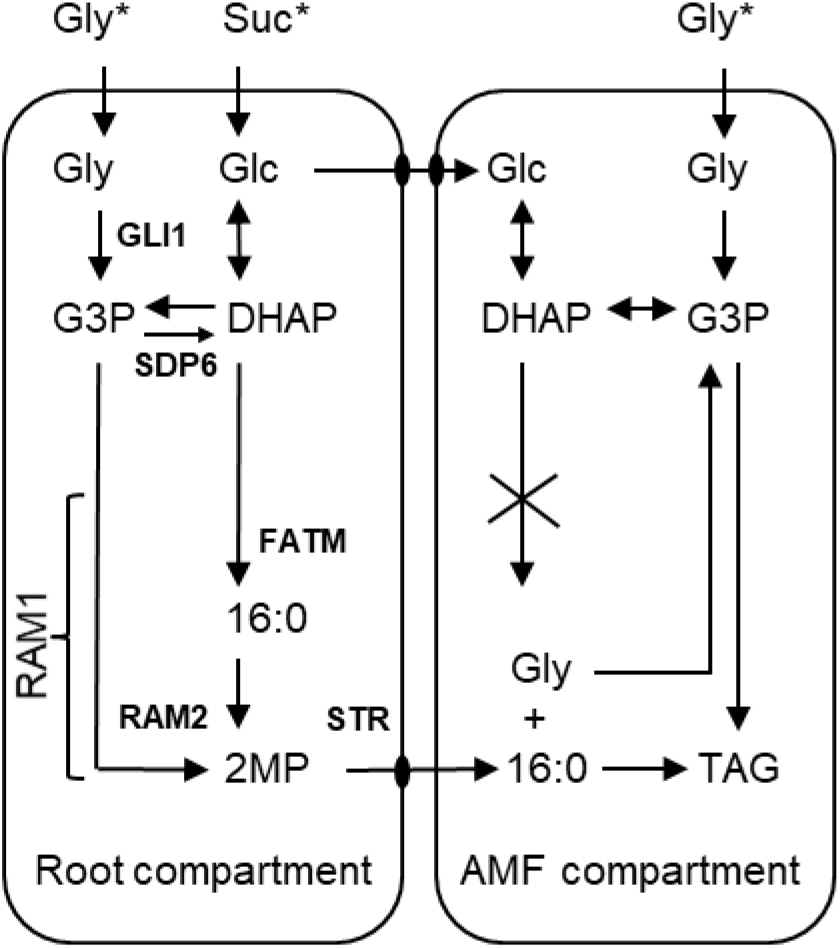
Diagram depicting model for plant lipid supply to AMF and metabolism of ^14^C glycerol and ^14^C sucrose. Asterisk denotes entry of ^14^C labelled substrate. Cross denotes deficiency in type-1 fatty acid synthase. Gly, glycerol; Glc, glucose; G3P, glycerol-3-phosphate; DHAP, dihydroxyacetonephosphate; 16:0 palmitic acid; 2MP, 2-monopalmitate; TAG, triacylglycerol; GLI1, glycerol kinase, SDP6, FAD-dependent G3P dehy drogenase; RAM2, sn-2 G3P acyltransferase/phosphatase; STR, ABC transporter; RAM1, GRAS-domain transcription factor.

Nutrient exchange during AMF symbiosis is mediated by specialised hyphal structures called arbuscules that form transiently within root cortical cells (arbuscocytes) of the host plant (Smith and Read, 2008). Biochemical and genetic studies suggest that plastidial FA biosynthesis is upregulated in arbuscocytes (Keymer et al., 2017) and is terminated by the acyl-acyl carrier protein thioesterase FATM to release palmitic acid (16:0) (Bravo et al., 2017; Brands et al., 2018; Fig. 1). This 16:0 then enters the cytosolic acyl-Coenzyme A (CoA) pool and is esterified to the sn-2 position of glycerol-3-phosphate (G3P) by the bifunctional acyltransferase/phosphatase REQUIRED FOR ARBUSCULAR MYCORRHIZATION 2 (RAM2), yielding 2-monopalmitin (2MP) (Wang et al., 2012; Luginbuehl et al., 2017; Fig. 1). It has been proposed that 2MP is then exported directly across the plant peri-arbuscular membrane (PAM), by the half-subunit ATP binding cassette transporters STUNTED ARBUSCULE (STR) and STR2 (Zhang et al., 2010; Bravo et al., 2017; Jiang et al., 2017; Keymer et al., 2017; Luginbuehl et al., 2017; Fig. 1). The FA and glyceryl moieties may then be taken up across the fungal plasma membrane either as 2MP or as free molecules following hydrolysis (Bravo et al., 2017; Fig. 1).

Plants can produce a wide range of extracellular lipids (Samuels et al., 2008; Yang et al., 2010) and fungi can assimilate many of them (Fickers et al., 2005). Although the existing pathway model (Fig. 1) is plausible, STR/STR2-dependent export of 2MP has yet to be demonstrated and the precise form(s) in which plants supply FA to AMF is still unclear. The metabolic tracer studies that provide direct evidence of plant lipid transfer to AMF were designed to follow the fate of FA and not glyceryl moieties (Jiang et al., 2017; Keymer et al., 2017; Luginbuehl et al., 2017). Interestingly, Jiang et al., (2017) observed that when ^13^C glycerol is supplied to mycorrhizal roots it labels AMF sugars more strongly than plant sugars. Assuming steady state labelling, this suggests a flux of ^13^C from glycerol to AMF sugars that bypasses plant sugars (Jiang et al., 2017). This could be due to transfer of glyceryl moieties in 2MP from plant to AMF, but it could also be explained by ^13^C glycerol labelling of plant FA and their subsequent transfer as 2MP, or by direct glycerol uptake and metabolism by AMF (Bago et al., 2002; Fig. 1). Jiang et al., (2017) also used 10 mM ^13^C glycerol, which is a high enough concentration to inhibit gluconeogenesis in plants (Aubert et al., 1994; Eastmond, 2004; Quettier et al., 2008). Interestingly, a recent study has also raised questions concerning the existing pathway model by showing that myristic acid (14:0) can support asymbiotic growth and metabolism of AMF, but neither 2MP nor 16:0 are able to function as effective carbon sources (Sugiura et al., 2020). The aim of this study was therefore to investigate whether the proposed pathway (Bravo et al., 2017; Fig. 1) can export 2MP from root cells, and whether AMF receive and utilise both the FA and glyceryl moieties.

## RESULTS

### *RAM1* expression drives 2MP secretion from *M. truncatula* roots

In *Medicago truncatula*, arbuscule development requires extensive transcriptional reprogramming of root cells mediated by a set of transcription factors that includes RAM1 (Gobbato et al., 2012; Park et al., 2015; Luginbuehl et al., 2017; Jiang et al., 2018). Ectopic expression of *RAM1* upregulates several components of the FA supply pathway in roots of *M. truncatula*; including *FATM, RAM2* (Luginbuehl et al., 2017) and *STR* (Park et al., 2015) (Fig. 1). We therefore decided to investigate whether *RAM1* could also trigger physiological expression of the pathway and drive 2MP export from *M. truncatula* roots. We transformed *M. truncatula* hairy roots with a construct expressing *RAM1* under the control of the constitutive *CaMV 35S* promoter (*Pro35S:RAM1*). We then grew the composite plants on synthetic growth medium, extracted nonpolar lipids from the surface of the roots by chloroform dipping and analysed them using gas chromatography (Li et al., 2007; Kosma et al., 2015). The main components of wild type *M. truncatula* root surface lipids are alkyl hydroxycinnamates and sterols, similar to other species that have been surveyed (Li et al., 2007; Kosma et al., 2015). However, the surfaces of roots expressing *RAM1* (Fig. 2A) exhibited a ∼30-fold increase in the abundance of monoacylglycerols (MGs) containing long-chain saturated FA moieties (Fig. 2B). 2MGs were the most abundant regioisomers, despite the fact they are more thermodynamically unstable (Li et al., 2007). The most abundant MG species contained 16:0 (i.e. 2MP), but species containing 14:0 and stearic acid (18:0) were also detected (Fig. 2B). No MGs with unsaturated FA moieties were detected. *RAM1* expression also led to a small increase in the abundance of 16:0, which is most likely to be a product of extracellular MG hydrolysis (Li et al., 2007). Our data suggest that *RAM1* expression can drive the production and secretion of 2MGs from *M. truncatula* roots, supporting the hypothesis that FA could predominantly be transported across the PAM in this esterified form.

**Fig. 2.**
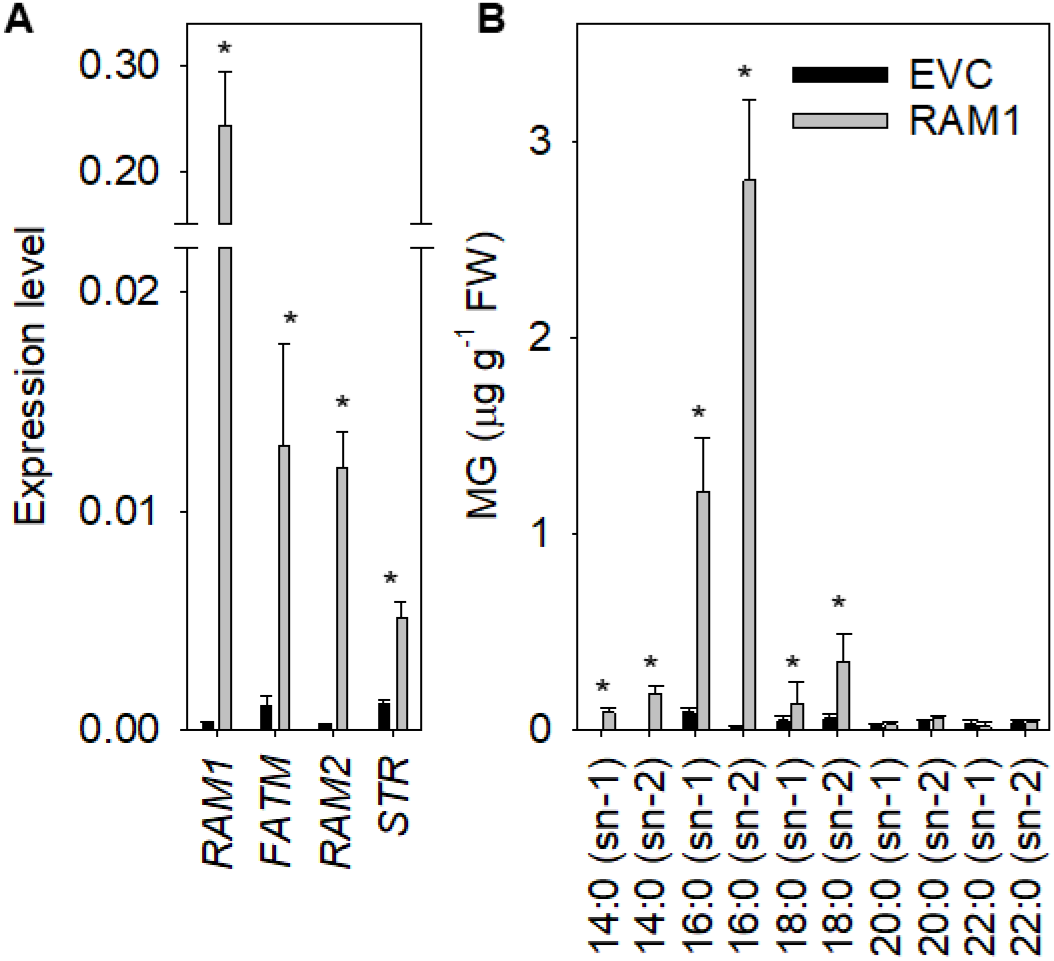
Expression of *RAM1* induces MG secretion from *M. truncatula* roots. (A) Q-PCR analysis of *RAM1, FATM, RAM2* and *STR* expression in roots ectopically expressing *RAM1*. (B) Effect of *RAM1* expression on MG content of root surface lipids. Values are the mean ± SE of measurements on five biological replicates. In A the expression is relative to the mean of the reference genes *UBI* and *EF1*. ^*^denotes values significantly (P < 0.05) different from EVC (ANOVA + Tukey HSD test).

### *RAM1*-dependent 2MP secretion requires *RAM2, FATM* and *STR*

It has previously been shown that one of RAM1’s direct transcriptional targets is *RAM2* (Gobbato et al., 2012). To determine whether RAM1-dependent MG secretion requires RAM2, we expressed *RAM1* in the roots of the *M. truncatula ram2-1* mutant (Wang et al., 2012) and analysed the surface lipids. We found that the accumulation of 2MP was virtually abolished in *ram2-1* (Fig. 3A). *FATM* and *STR* are also induced by RAM1 (Luginbuehl et al., 2017; Park et al., 2015; Fig 2A). To investigate whether FATM and STR are necessary for RAM1-mediated MG secretion, we designed two RNA interference (RNAi) constructs to silence the expression of each of these target genes (Zhang et al., 2010; Jiang et al., 2017) and added a *Pro35S:RAM1* cassette to simultaneously drive *RAM1* expression. In control experiments we showed that the RNAi constructs were able to supress RAM1-dependent expression of their target genes, when they were transformed into *M. truncatula* hairy roots, and to inhibit mycorrhization when the roots were colonised with the AMF *Rhizophagus irregularis* (Table S1). When we transformed these constructs into uncolonized hairy roots, we found that they also supressed RAM1-dependent secretion of 2MP (Fig. 3B). These data show that *RAM2, FATM* and *STR* are all required for RAM1-mediated MG secretion and this finding is consistent with a pathway model in which STR/STR2 transport 2MGs across the PAM (Bravo et al., 2017).

**Fig 3.**
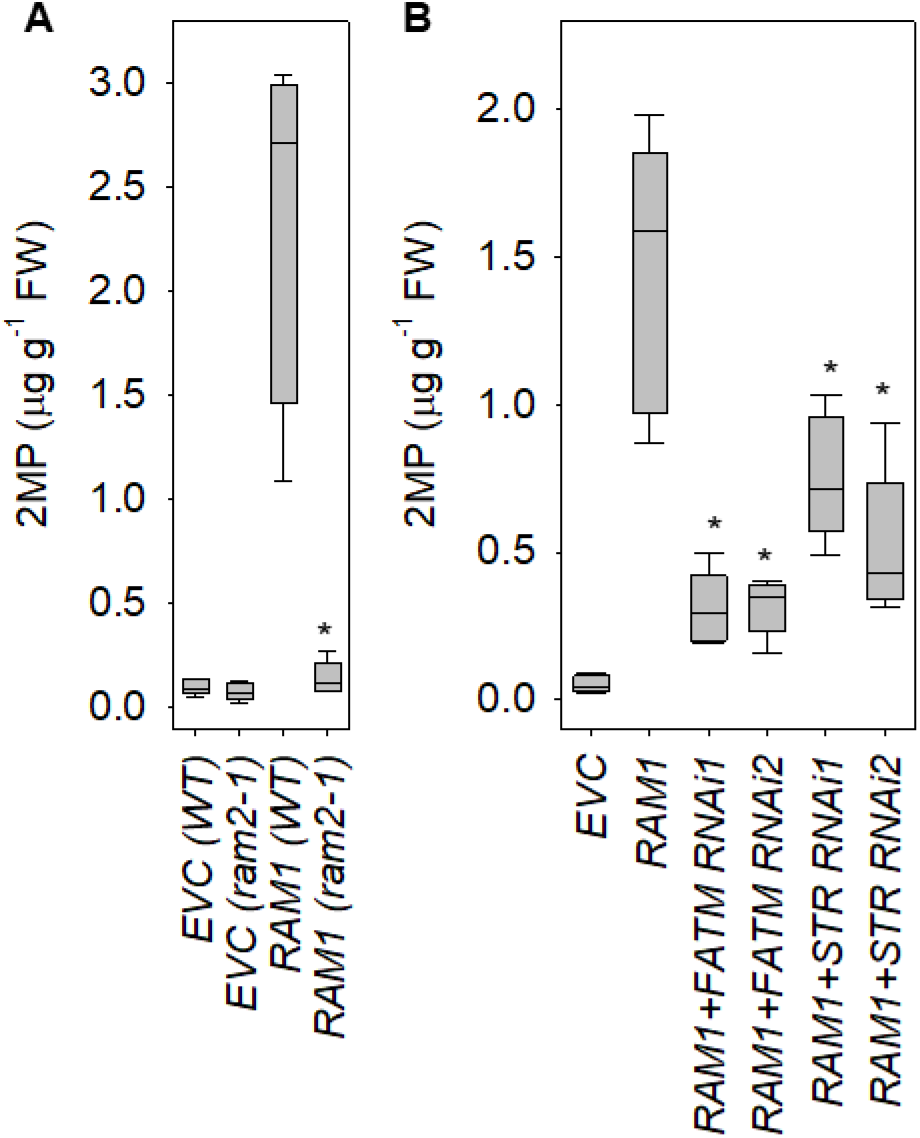
RAM1-dependent 2MP secretion requires RAM2, FATM and STR. (A) Effect of *RAM1* expression in *ram2-1* roots. (B) Effect of *RAM1* expression in *FATM RNAi* and *STR RNAi* roots. Values are the mean ± SE of measurements on five biological replicates. ^*^denotes values significantly (P < 0.05) different from *RAM1* (ANOVA + Tukey HSD test).

### GLI1 is required for ^14^C glycerol labelling of FA and glyceryl moieties of AMF lipids

Isotopically labelled acetate and glycerol are routinely used to trace the flux of FA and glyceryl moieties in plant lipid metabolism (Allen et al., 2015). Neither metabolite is a direct physiological precursor for glycerolipid biosynthesis in plants, but acetate is converted to acetyl-CoA by plastidial acetyl-CoA synthetase (ACS) (Lin and Oliver, 2008) and glycerol to G3P by the glycerol kinase GLYCEROL INSENSITIVE 1 (GLI1) (Eastmond, 2004; Fig. 1). We previously performed ^14^C acetate labelling experiments on mycorrhizal roots of a *M. truncatula acs* mutant to show that the FA moieties in AMF TAG are supplied by their host plant (Luginbuehl et al., 2017). To determine whether the glyceryl moieties also come from the host plant we identified a *M. truncatula gli1* mutant (Fig. 4). The *M. truncatula* genome contains a single *GLI1* homologue (Medtr4g009920) that shares 78% amino acid identity with its *A. thaliana* counterpart (Eastmond, 2004) (Fig. S1) and is expressed in roots based on RNA-seq data (Luginbuehl et al., 2017). A *Tnt1* retrotransposon tagged (Tadege et al., 2008) mutant of *GLI1* was identified by PCR-based reverse screening (Cheng et al., 2014) and the insertion site in the first intron was confirmed by amplifying and sequencing genomic PCR products that span the *Tnt1* borders (Fig. 4A). In contrast to wild type, homozygous *gli1* roots were found to lack *GLI1* expression and glycerol kinase activity and were unable to metabolise exogenously applied 0.5 mM [U-^14^C] glycerol to lipids (Fig. 4B-D). However, glycerol kinase activity and ^14^C glycerol labelling of lipids were both rescued by hairy root transformation of *gli1* with a *Pro35S:GLI1* construct (Fig. 4B-D). These data showed that GLI1 is required for glycerol metabolism in *M. truncatula* roots.

**Fig. 4.**
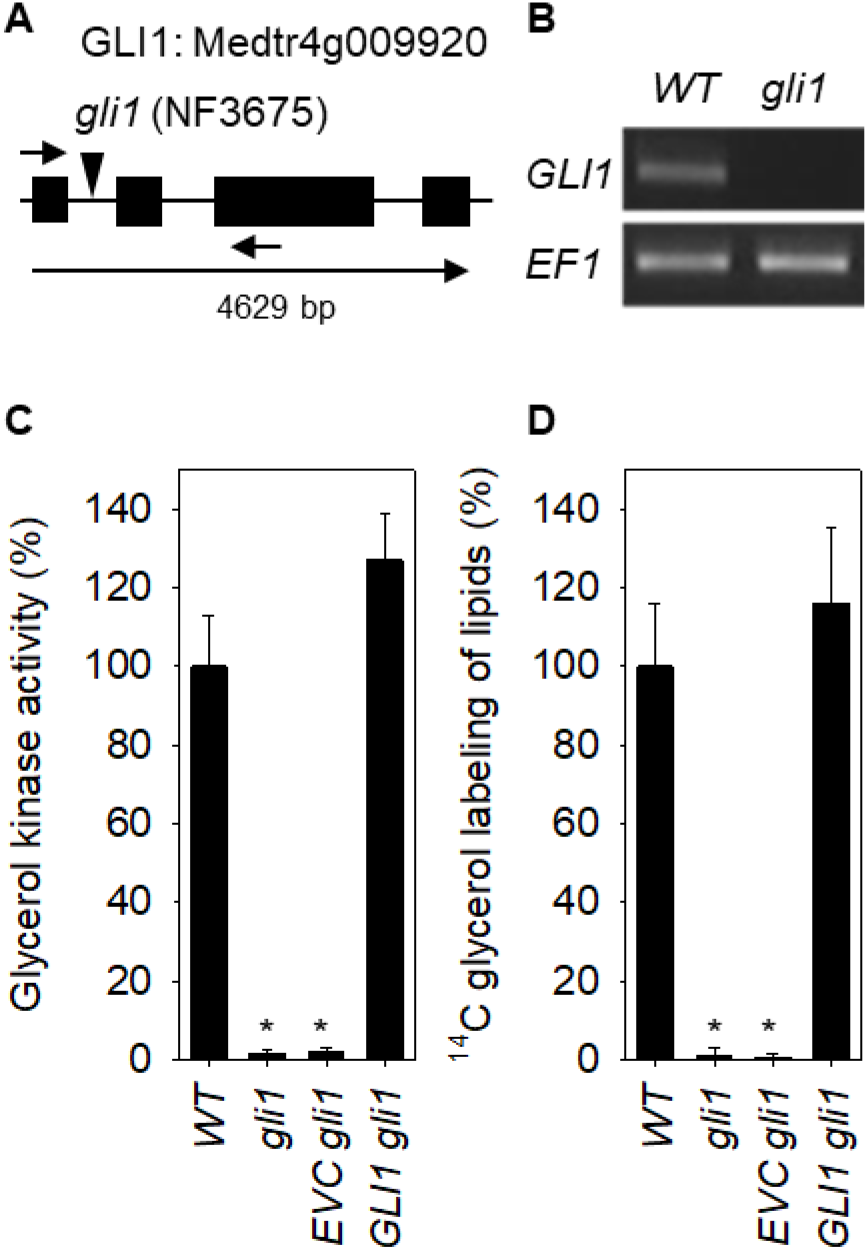
GLI1 is required for glycerol metabolism in *M. truncatula* roots. (A) Model of *GLI1* showing position of *Tnt1* transposon insertion. (B) RT-PCR analysis of *GLI1* expression in *gli1* roots. (C) Glycerol kinase activity in *gli1* roots. (D) Incorporation of ^14^C glycerol into *gli1* root lipids. Values are the mean ± SE of measurements on five root systems and are expressed as a percentage of WT. Values are the mean ± SE of measurements on five biological replicates. *denotes values significantly (P < 0.05) different from WT (ANOVA + Tukey HSD test).

The *M. truncatula gli1* mutant lacked an obvious phenotype under normal growth conditions. However, like *A. thaliana gli1* (Eastmond, 2004), seedling root growth is inhibited by high concentrations of glycerol (Table S2), that lie outside of the 0.1 to 0.5 mM physiological range (Gerber et al., 1988). We colonised *gli1* roots with *R. irregularis*, fed them with 0.5 mM [U-^14^C] glycerol or [U-^14^C] sucrose and measured ^14^C in both the FA and glyceryl moieties of TAG (Fig. 5). Virtually all the TAG is of AMF origin because roots colonised with *R. irregularis* contain around 70 times more TAG than uncolonized roots and the unique AMF marker FA palmitvaccenic acid (16:1^Δ11cis^) (Trépanier et al., 2005) is esterified to the middle (sn-2) position on ∼95% of the glyceryl moieties (Table S3). In mycorrhizal wild type roots, ^14^C from glycerol was readily incorporated into TAG and only around 6% of the ^14^C was found in the FA moieties (Fig. 5B&C). In mycorrhizal *gli1* roots, we found that the incorporation of ^14^C from glycerol into TAG was reduced by around 80% and no ^14^C was present in the FA moieties (Fig. 5B&C). In contrast, the incorporation of label from 0.5 mM [U-^14^C] sucrose into TAG was not impaired in mycorrhizal *gli1* roots (Fig. 5D) and the percentage of ^14^C found in the FA moieties was around 92% (Fig. 5E). These data suggest that most of the glyceryl moieties (and all the FA moieties) in AMF TAG are derived from ^14^C glycerol metabolism in the host plant and not in the AMF (Fig. 1). Furthermore, the preferential labelling of the glyceryl moiety of TAG by ^14^C glycerol implies that this flux does not proceed via sugars (Fig. 1). Our ^14^C sucrose labelling data suggest that, if this were the case then, a far higher proportion of ^14^C should be found in the FA moieties.

**Fig. 5.**
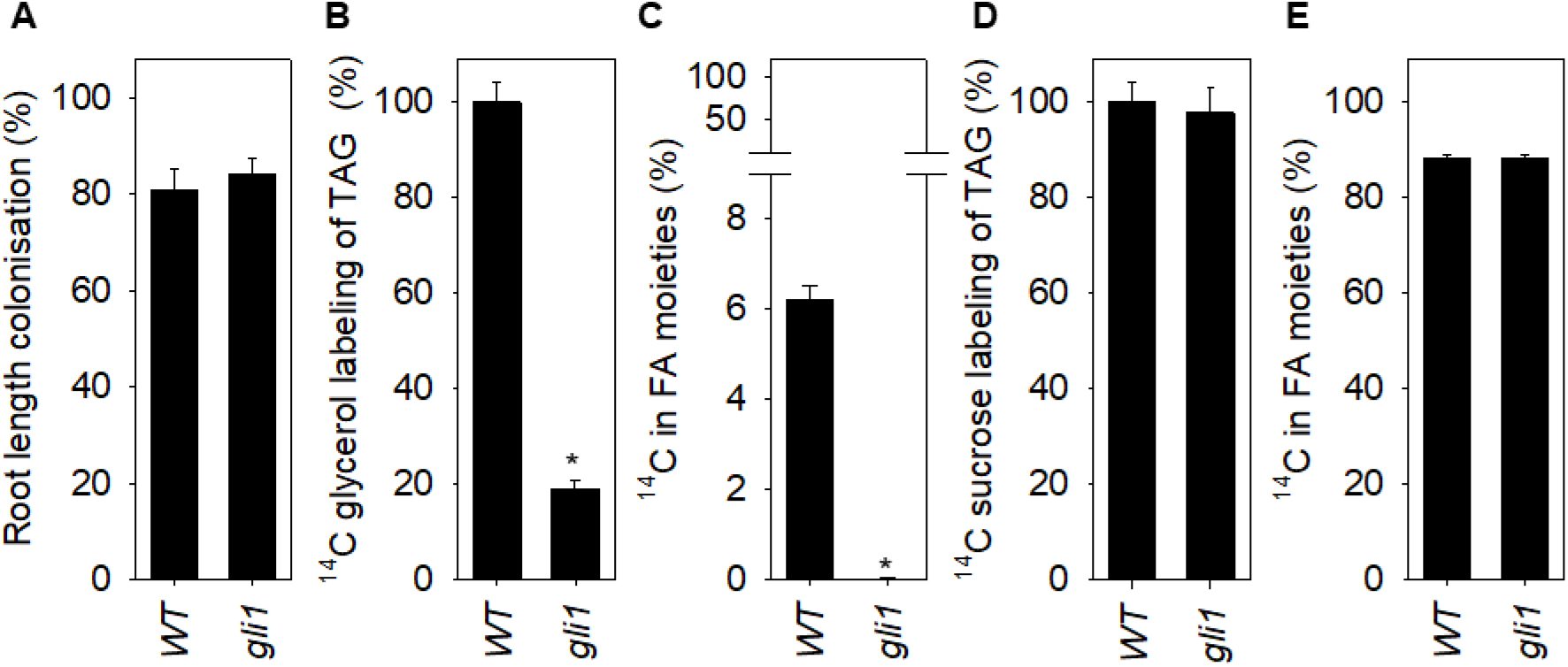
GLI1 is required for ^14^C glycerol labelling of FA and glycerol moieties of AMF TAG. (A) Root length colonisation by *R. irregularis* at five weeks post inoculation. Incorporation of ^14^C glycerol (B) and ^14^C sucrose (D) into TAG from mycorrhizal roots. Percentage of ^14^C in FA moiety of TAG, respectively (C and E). Values are the mean ± SE of measurements on five biological replicates and are expressed as a percentage of WT in B and D. ^*^denotes values significantly (P < 0.05) different from WT (two-tailed Student’s t test).

### SDP6 is only required for ^14^C glycerol labelling of the FA moieties of AMF lipids

Labelling of FA by ^14^C glycerol is made possible by metabolism of G3P to the glycolytic intermediate dihydroxyacetonephosphate (DHAP) (Slack et al., 1977; Bates et al., 2008; Fig. 1). This reaction is mainly catalysed by the mitochondrial FAD-dependent G3P dehydrogenase (FAD-G3PDH) SUGAR-DEPENDENT6 (SDP6) (Quettier et al., 2008), while the small equilibrium constant of NAD-dependent G3P dehydrogenase strongly favours the reverse reaction (Quettier et al., 2008). The *M. truncatula* genome contains two SDP6 homologues (SDP6a; Medtr1g094185 and SDP6b; Medtr1g050720) that share 78% amino acid identity (outside of their N-terminal mitochondrial targeting sequence) with their *A. thaliana* counterpart (Quettier et al., 2008) (Fig. S2) and 84% nucleotide identity with each other. RNA-seq data suggest that both *SDP6* genes are expressed in roots (Luginbuehl et al., 2017). To investigate whether SDP6 is necessary for ^14^C glycerol labelling of AMF TAG, we designed two RNA interference (RNAi) constructs to silence the expression of both *SDP6* homologues (Zhang et al., 2010; Jiang et al., 2017). In control experiments we showed that the constructs were able to supress the transcript abundance of their target genes and reduce FAD-G3PDH activity, when they were transformed into wild type *M. truncatula* hairy roots (Table S4). When roots transformed with the SDP6 RNAi constructs were colonised with *R. irregularis* (Fig. 6A) and fed with 0.5 mM [U-^14^C] glycerol, labelling of TAG was not significantly affected (Fig. 6B), but the percentage of ^14^C found in FA moieties was reduced to less than 1% (Fig. 6C). These data confirm that ^14^C glycerol does not label the glyceryl moiety of AMF TAG via plant DHAP and downstream metabolites such as sugars (Fig. 1).

**Fig. 6.**
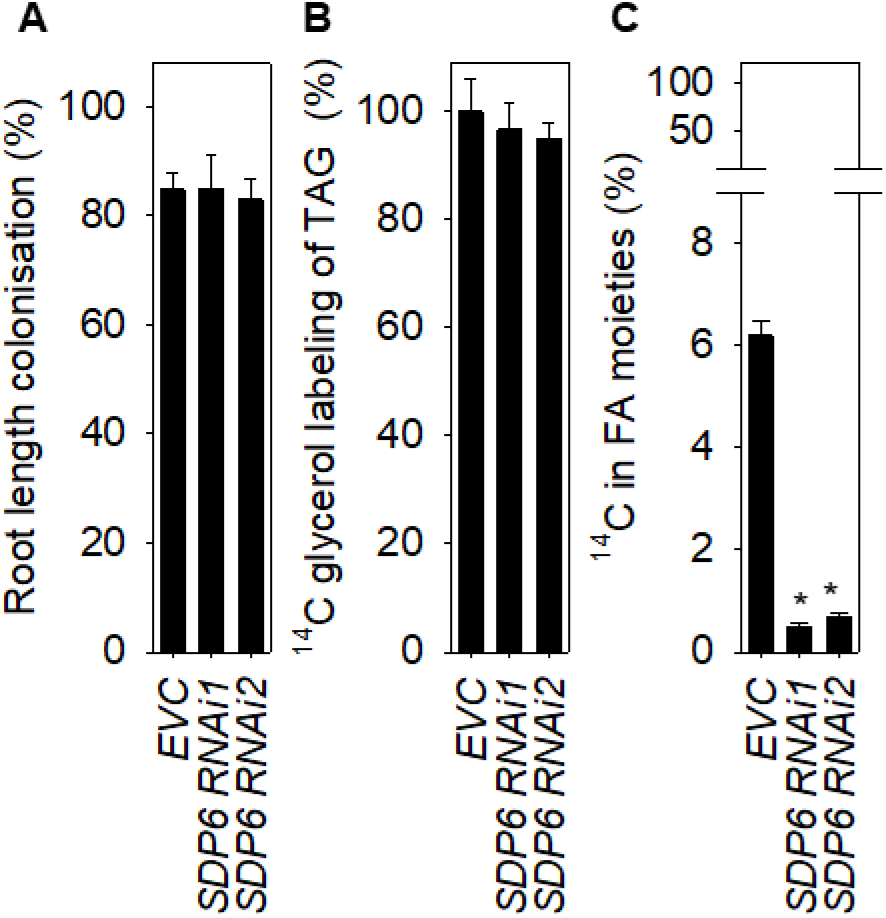
SDP6 is required for ^14^C glycerol labelling of FA but not glyceryl moieties of AMF TAG. (A) Root length colonization by *R. irregularis* at five weeks post inoculation. (B) Incorporation of ^14^C glycerol into TAG from mycorrhizal roots. (C) Percentage of ^14^C in the FA moiety of TAG. Values are the mean ± SE of measurements on five biological replicates and are expressed as a percentage of the empty vector control (EVC) in B. ^*^denotes values significantly (P < 0.05) different from EVC (ANOVA + Tukey HSD test).

### RAM2 is required for ^14^C glycerol labelling of FA and glyceryl moieties of AMF lipids

The remaining route for G3P metabolism in plants is via glycerolipid biosynthesis (Slack et al., 1977; Bates et al., 2008). The supply of plant FA moieties to AMF is known to rely on the G3P acyltransferase RAM2 (Jiang et al., 2017; Luginbuehl et al., 2017; Keymer et al., 2017). Therefore, it is most probable that the glyceryl moieties of AMF TAG are supplied via RAM2-mediated acylation of G3P (Fig. 1). To investigate whether RAM2 is involved in the provision of both FA and glyceryl moieties present in AMF TAG, we performed ^14^C glycerol labelling experiments on heterozygous *ram2*/*RAM2* roots colonised with *R. irregularis*. Although mycorrhization is severely compromised in *ram2* roots (Wang et al., 2012) and the arbuscules are stunted/collapsed (Bravo et al., 2007), we found that roots length colonisation (Fig. 7A), the percentage of root segments with WT arbuscules and mean arbuscule length (Table S5 and Fig. S3) were not reduced in *ram2*/*RAM2*. Zhang et al., (2010) also reported that *str* may be recessive. The expression level of *PHOSPHATE TRANSPORTER 4* (*PT4*) was also not reduced in mycorrhizal *ram2/RAM2* (Fig. 7B). *PT4* is specifically expressed in arbuscocytes, under the control of RAM1 (Luginbuehl et al., 2017) and encodes a PAM protein that is required for phosphate uptake (Javot et al., 2007). Unlike *PT4*, the expression of *RAM2* was reduced in *ram2/RAM2* by nearly 50% (Fig. 7C). Incorporation of ^14^C glycerol into TAG was also reduced by around 25% in *ram2*/*RAM2* roots (Fig. 7C) and the percentage of ^14^C in the FA moieties remained at around 6% (Fig. 7D). These data suggest that the supply of both FA and glyceryl moieties of AMF TAG relies on the level of *RAM2* expression and RAM2 may exert a significant degree of control over this metabolic flux.

**Fig. 7.**
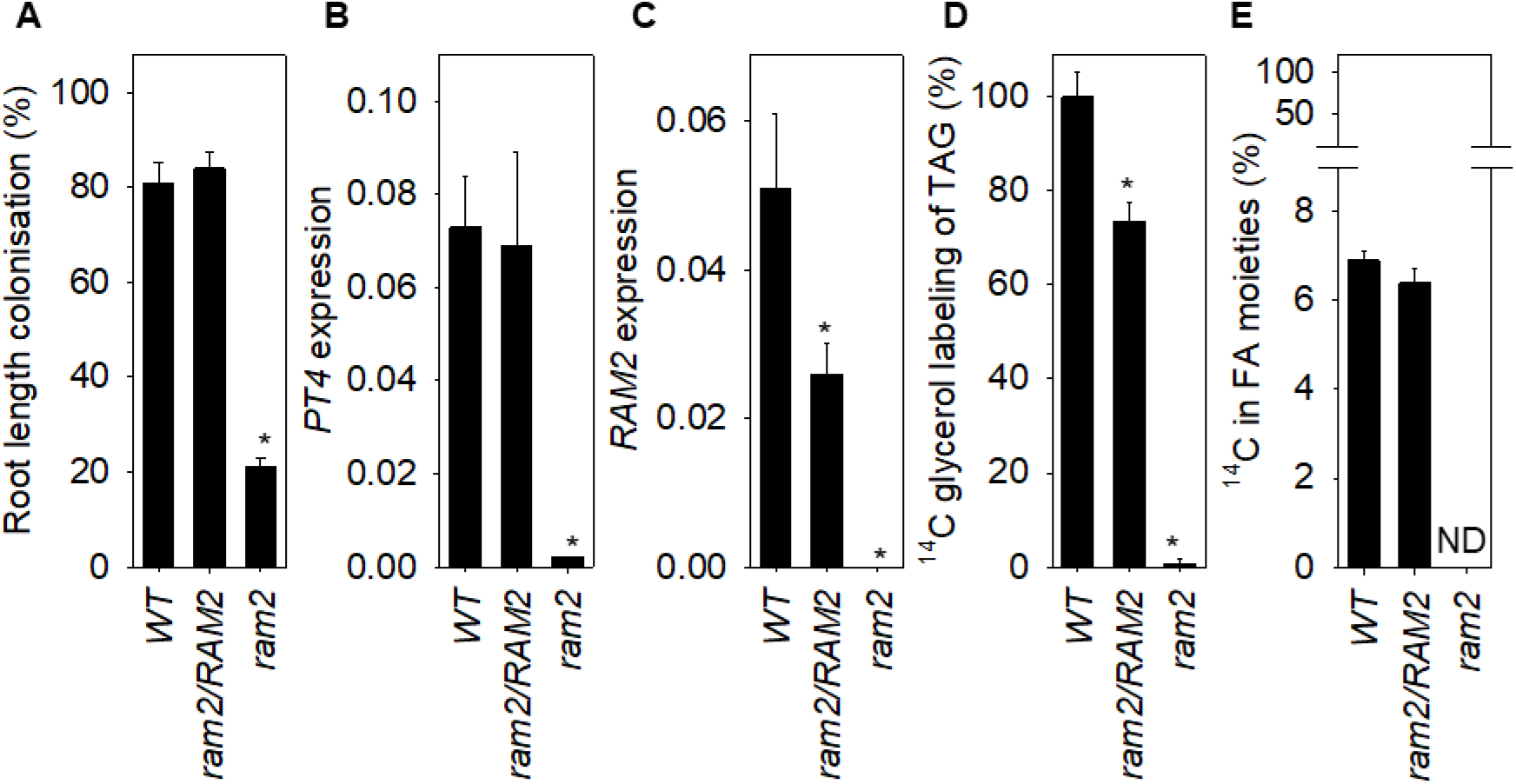
*RAM2* is required for ^14^C glycerol labelling of both moieties of AMF TAG. (A) Root length colonisation by *R. irregularis* five weeks post inoculation. Q-PCR analysis of *PT4* (B) and *RAM2* expression. (D) Incorporation of ^14^C glycerol into TG from mycorrhizal roots. (E) Percentage of ^14^C in FA moieties of TAG. In B and C, expression is relative to the mean of the reference genes *UBI* and *EF1*. Values are the mean ± SE of measurements on five biological replicates and are expressed as a percentage of WT in D. ^*^denotes values significantly (P < 0.05) different from WT (ANOVA + Tukey HSD test). ND is not determined.

## DISCUSSION

In this study we show that the RAM1-dependent transcriptional program that accommodates arbuscules can induce 2MG secretion from roots that relies on FATM, RAM2 and STR, and that AMF receive both FA and glyceryl moieties from their host plant and use them to make TAG. The predominant form of 2MG that is secreted from roots when RAM1 is ectopically expressed is 2MP, which is consistent with the proposed model for plant lipid transfer to AMF (Bravo et al., 2017; Fig. 1). However, low levels of 2MG species containing 14:0 and 18:0 were also detected. This is consistent with the *in vitro* substrate specificities of FATM (Brands et al., 2018) and RAM2 (Luginbuehl et al., 2017). Asymbiotic culture experiments also suggest that 14:0 is necessary to sustain AMF and this may be due to a requirement for myristoylation (Sugiura et al., 2012). The dependence of 2MG export on STR also supports the hypothesis that STR/STR2 transports this substrate across the PAM (Bravo et al., 2017). However, *in vitro* characterisation of purified STR/STR2 is necessary to ascribe this biochemical function (Gräfe et al., 2020).

The *M. truncatula* glycerol kinase gene *GLI1* is necessary for mycorrhizal roots to incorporate label from ^14^C glycerol into both the FA and glyceryl moieties of *R. irregularis* TAG. Although *R. irregularis* can take up and metabolise glycerol directly (Bago et al., 2002; Fig.1), our data show that more than 80% of the glycerol is incorporated into the glyceryl moiety of AMF TAG via root metabolism (Fig. 1). There is already substantial evidence that roots provide essentially all the FA moieties present in AMF lipids and that these moieties are supplied by a 2MG pathway that utilises the sn-2 G3P acyltransferase/phosphatase RAM2 (Jiang et al., 2017; Luginbuehl et al., 2017; Keymer et al., 2017). Acylation of G3P by RAM2, followed by export of both 2MG moieties to the AMF would therefore account for their GLI1-dependent labelling by ^14^C glycerol (Fig. 1).

The alternate explanation is that glycerol is metabolised within the root to other transferable intermediates such as sugars (Shachar-Hill et al., 1995; Solaiman and Saito, 1997) or FA (Jiang et al., 2017; Keymer et al., 2017; Luginbuehl et al., 2017) that are then taken up by the AMF and converted to G3P. Although these metabolic routes are feasible, they cannot account for the ^14^C glycerol labelling pattern of AMF TAG, in which ∼94% of the ^14^C is found in the glyceryl moiety. This strong preferential labelling of the glyceryl moiety is common in plant tissues (Slack et al., 1977; Bates et al., 2008) and has also been observed previously in AMF TAG, when mycorrhizal roots are labelled with ^13^C glycerol (Bago et al., 2002). In plant tissues, the labelling imbalance between the glyceryl and FA moieties likely reflects the high rate of glycerol metabolism to G3P relative to DHAP (Aubert et al., 1994; Eastmond, 2004; Quettier et al., 2008). Supplying mycorrhizal roots with ^14^C sucrose (this study), or ^13^C glucose (Pfeffer et al., 1999), results in a much higher proportion of label in the FA moieties of AMF TAG and this would also be expected if ^14^C glycerol labelled the FA and glyceryl moieties of AMF TAG via any intermediates down-stream of DHAP, such as sugars or FA (Fig. 1). G3P is mainly converted to DHAP by FAD-G3PDH in *A. thaliana* (Quettier et al., 2008). We found that RNAi suppression of the FAD-G3PDH genes *SDP6a* and *SDP6b* in *M. truncatula* virtually abolished ^14^C glycerol labelling of the FA moiety of AMF TAG, which confirms that labelling of the glyceryl moiety does not proceed via DHAP (Fig. 1).

Further evidence that both FA and glyceryl moieties of AMF TAG are supplied via the 2MG pathway was obtained by performing ^14^C glycerol labelling experiments on mycorrhizal roots of *ram2*/*RAM2* plants. Homozygous *ram2* roots are severely impaired in mycorrhization (Wang et al., 2012; Bravo et al., 2017), rendering the interpretation of labelling experiments problematic (Keymer et al., 2017). However, *ram2*/*RAM2* roots colonised with *R. irregularis* were indistinguishable from wild type, suggesting that *ram2* is recessive in terms of its gross mycorrhization phenotype. Despite this, heterozygous mutants in plant enzymes often exhibit a fractional reduction in gene expression and activity that can be used to study effects on metabolic flux (Wingler et al., 2002). *RAM2* expression was reduced by nearly 50% and ^14^C glycerol labelling of AMF TAG by ∼25% for both FA and glyceryl moieties. The provision of both moieties of AMF TAG is therefore equally reliant on *RAM2*. The responsiveness of flux to a fractional reduction in *RAM2* expression also suggest that RAM2 activity might exert significant control over the pathway (Wingler et al., 2002). Li et al., (2007) also showed that over-expression of the *A. thaliana* sn-2 G3P acyltransferase GPAT5, which is required for biosynthesis of suberin monomers, can enhance 2MG secretion.

Taken together, data from ectopic RAM1 expression and labelling studies using *gli1, SDP6 RNAi* and *ram2*/*RAM2* are consistent with a pathway model in which both moieties of the 2MGs produced by RAM2 are supplied to AMF (Fig. 1). However, it is not known whether 2MGs traverse the fungal plasma membrane, intact or are hydrolysed first. The latter is more probable given that FA and not 2MGs have recently been shown to support asymbiotic growth and metabolism of AMF (Sugiura et al., 2012). Fungi commonly secrete extracellular lipases to allow assimilation of FA esters (Fickers et al., 2005) and this may be necessary within the symbiotic phase of the AMF life cycle. It is also likely that AMF assemble their glycerolipids from FA and glycerol (Fig. 1) rather than from 2MGs, given that the 2MGs appear to comprise mainly 16:0 moieties and ∼95% of the FA moieties at the sn-2 position in *R. irregularis* TAG are 16:1^Δ11cis^ (Table S3), which is most likely produced by a fungal fatty acyl-CoA Δ11 desaturase (Cheeld et al., 2020; Brands et al., 2020). Surprisingly, 16:0 cannot support asymbiotic growth of AMF, although it can be taken up, desaturated to 16:1^Δ11cis^ and incorporated into TAG (Sugiura et al., 2020). TAG is the major storage reserve in AMF spores and its dissimilation is presumed to contribute to asymbiotic growth of AMF (Bago et al., 2002). This apparent contradiction might be explained by channelling of 16:0 into storage lipids rather than β-oxidation. Although Sugiura et al., (2020) showed that 14:0 can support asymbiotic growth of AMF, ectopic expression of RAM1 suggests that 14:0 is only a minor component of the plant lipids exported to AMF during symbiosis. Keymer et al., (2017) also reported that the isotopolog profiles of plant and AMF 16:0 are identical and this is inconsistent with a pathway model where plant 14:0 is elongated to 16:0 by the fungus.

## METHODS

### Plant material and growth conditions

The *M. truncatula* R108 mutant *gli1* (NF3675) was identified by PCR-based reverse screening (Cheng et al., 2014) of the *Tnt1* tagged lines (Tadege et al., 2008) at the Noble Research Institute. The genotype of the mutant was confirmed by performing PCR on genomic DNA (Luginbuehl et al., 2017) using primers listed in Table S6. The *M. truncatula* Jemalong A17 mutant *ram2-1* has been described previously (Wang et al., 2012). Seeds were scarified, surface sterilized with 10% (v/v) bleach solution and plated on agar plates containing one-half strength Murashige and Skoog (MS) salts (Merck) pH 5.7. For mycorrhization experiments, seedlings were transplanted into a sterile mixture of terragreen and sand (1:1 v/v) and inoculated with *R. irregularis* (Luginbuehl et al., 2017). For root surface lipid analysis, seedlings were transplanted onto vertically orientated plates containing filter paper soaked in one-half strength MS salts plus 1% (w/v) sucrose pH 5.7. Plants were grown in a controlled environment chamber set to a 16 h light (22°C) / 8 h dark (18°C) period (photon flux density = 250 µmol m^2^ s^-1^).

### Gene expression analysis and enzyme assays

RNA was extracted from ∼100 mg of root tissue and DNase treated using the RNeasy plant mini kit and RNase free DNase kit (Qiagen) according to the manufacturer’s instructions. Reverse transcription was carried out with 1 μg of RNA using the SuperScript IV First-Strand Synthesis System (ThermoFisher Scientific) according to the manufacturer’s instructions. Quantitative PCRs were performed as described previously (Luginbuehl et al., 2017), using the 2^-ΔΔCt^ method (Livak and Schmittgen, 2001) and both *UBIQUITIN* (*UBI*) and *ELONGATION FACTOR 1 ALPHA* (*EF1*) as reference genes. The primer pairs used for gene expression analysis are listed in Table S6. Glycerol kinase and FAD-dependent G3P dehydrogenase assays were performed as described previously (Eastmond, 2004; Quettier et al., 2008).

### Cloning and hairy root transformation

For overexpression studies, *RAM1* and *GLI1* were amplified from root cDNA using the primer pair listed in Table S6. The products were cloned using the pENTR/D-TOPO Cloning Kit and transferred into pK7WG2R using Gateway LR reactions (Invitrogen). For RNAi studies, selected regions of *FATM, STR* and *SDP6a* were amplified from root cDNA using the primer pair listed in Table S6. The products were cloned using a pENTR/D-TOPO Cloning Kit and transferred into pK7GWIWG2(II)-RedRoot using Gateway LR reactions (Invitrogen). The *35S:RAM1* cassette from pK7WG2R was amplified using the primer pair listed in Table S1 and cloned into pK7GWIWG2(II)-RedRoot at the Kpn1 and Xho1 sites using T4 DNA ligase (NEB). The vectors were transformed into *M. truncatula* roots using *Agrobacterium rhizogenes* strain AR1193, following the protocol described previously (Boisson-Dernier et al., 2001). After transformation, 100 mg L^-1^ cefotaxime was added to the growth medium to supress *A. rhizogenes* growth. Untransformed roots not expressing the DsRed reporter were removed after three weeks and the composite plants were used for experiments.

### Staining, microscopy and quantification of colonisation

Mycorrhizal roots were washed with water, treated with 10% (w/v) KOH for 6 min at 95°C and stained with black ink for 3 min (Vierheilig et al., 1998). Mycorrhizal roots were also stained using WGA-AlexaFluor 488 (Molecular probes) and Nile Red (Merck) as described previously (Luginbuehl et al., 2017). Roots were imaged with a Zeiss LSM780 microscope using standard settings for WGA-AlexaFluor 488 and Nile Red. Root length colonization was quantified using the grid line intersect method described by McGonigle et al., (1990) and colonised segments were scored for WT arbuscules (Bravo et al., 2017).

### Root surface lipid analysis

Root surface lipids were extracted and analysed as described by (Li et al., 2007), with the following modifications. *M. truncatula* hairy roots were removed from plates, air dried at 50°C for 20 min and dipped in chloroform for 1 min. The extracts were filtered through glass wool, evaporated to dryness under a stream of N_2_gas and then derivatized by heating at 110°C for 10 min in pyridine:N,O-bis(trimethylsilyl)-trifluoroacetamide (1:1 [v/v]). The silylated samples were analyzed by gas chromatography (GC) using a 30 m HP-1ms capillary column (Agilent). For molecular identification, a Hewlett-Packard 5890 GC-coupled MSD 5972 mass analyzer was used with the mass analyzer set in electron impact mode (70 eV) and scanning from 40 to 700 atomic mass units. MG isomers were distinguished both by retention time and by their diagnostic fragmentation patterns (Li et al., 2007; Destaillats et al., 2010). A Hewlett-Packard 5890 GC with flame ionization detector was used for quantification, together with MG standards (Merck). Due to complex root architecture, it was not practical to calculate lipid content based on surface area and we report lipids as micrograms per gram fresh weight.

### ^14^C labelling of root lipids

Wild-type, mutant and hairy root-transformed plants were grown in a sterile mixture of terra green and sand (1:1 v/v) inoculated with *R. irregularis* and they were watered every two days. 5 μCi of [U-^14^C] glycerol or [U-^14^C] sucrose (PerkinElmer) at a concentration of 0.5 mM were added to the growth medium five weeks after inoculation. Two weeks later the colonized root systems were harvested, a tripentadecanoin standard (Merck) was added and total lipids were extracted and the triacylglycerol (TAG) was purified by thin layer chromatography (Kelly et al., 2013). Fatty acid methyl esters (FAMEs) were prepared from the TAG by transesterification and their quantity and molecular identity was determined by GC (Kelly et al., 2013). Fatty acyl composition at the sn-2 position of TAG was also determined, as described previously (van Erp et al., 2019). The ^14^C content of the FAMEs and glycerol backbone (Bates et al., 2008) were measured by liquid scintillation counting and their specific activities were calculated.

### Statistical Analyses

All experiments were carried out using five biological replicates and the data are presented as the mean values ± standard error of the mean (SE). For statistical analysis we either used one-way analysis of variance (ANOVA) with post-hoc Tukey HSD (Honestly Significant Difference) tests, or two-tailed Student’s t-tests.

### Accession numbers

The gene identifiers for the sequences used in this study are as follows: Medtr7g027190 (RAM1), Medtr1g040500 (RAM2), Medtr1g109110 (FATM), Medtr8g107450 (STR), Medtr4g009920 (GLI1), Medtr1g094185 (SDP6a), Medtr1g050720 (SDP6b), Medtr1g028600 (PT4).

## Supplemental data

Supplemental Figure 1. Amino acid sequence alignment of *A. thaliana* and *M. truncatula* GLI1.

Supplemental Figure 2. Amino acid sequence alignment of *A. thaliana* and *M. truncatula* SDP6.

Supplemental Figure 3. Arbuscules in heterozygous *ram2* roots colonised by *R. irregularis*.

Supplemental Table 1. Effect of *FATM* and *STR RNAi* on RAM1-dependent expression and mycorrhization.

Supplemental Table 2. Effect of exogenous glycerol on *gli1* seedling root growth. Supplemental Table 3. TAG composition of *M. trucatula* roots colonised with *R. irregularis*.

Supplemental Table 4. Effect of *SDP6 RNAi* on *SDP6* expression and FAD-G3PDH activity.

Supplemental Table 5. Colonisation of heterozygous *ram2* roots by *R. irregularis*. Supplemental Table 6. PCR primers used

## Acknowledgments

We thank Dr Frederick Beaudoin for providing his expertise and assistance in GC-MS analysis. This work was funded by the UK Biotechnology and Biological Sciences Research Council through grants BB/P012663/1, BB/J004553/1 and BB/K003712/1. *M. truncatula Tnt1* mutants were created through research funded, in part, by a grant from the National Science Foundation, DBI-0703285 and IOS-1127155.

## Author contributions

P.J.E and G.E.D.O. designed research; L.H.L., H.vE. H.C. and P.J.E. performed research; K.S.M and J.W. provided reagents; L.H.L., H.vE. and P.J.E. analyzed data; and P.J.E. wrote the paper, with contributions from all authors.

